# Efficacy of late-onset antiviral treatment in immune-compromised hosts with persistent SARS-CoV-2 infection

**DOI:** 10.1101/2024.05.23.595478

**Authors:** Carolin M Lieber, Hae-Ji Kang, Elizabeth B Sobolik, Zachary M Sticher, Vu L Ngo, Andrew T Gewirtz, Alexander A Kolykhalov, Michael G Natchus, Alexander L Greninger, Mehul S Suthar, Richard K Plemper

## Abstract

The immunocompromised are at high risk of prolonged SARS-CoV-2 infection and progression to severe COVID-19. However, efficacy of late-onset direct-acting antiviral (DAA) therapy with therapeutics in clinical use and experimental drugs to mitigate persistent viral replication is unclear. In this study, we employed an immunocompromised mouse model, which supports prolonged replication of SARS-CoV-2 to explore late-onset treatment options. Tandem immuno-depletion of CD4^+^ and CD8^+^ T cells in C57BL/6 mice followed by infection with SARS-CoV-2 variant of concern (VOC) beta B.1.351 resulted in prolonged infection with virus replication for five weeks after inoculation. Early-onset treatment with nirmatrelvir/ritonavir (paxlovid) or molnupiravir was only moderately efficacious, whereas the experimental therapeutic 4’-fluorourdine (4’-FlU, EIDD-2749) significantly reduced virus load in upper and lower respiratory compartments four days post infection (dpi). All antivirals significantly lowered virus burden in a 7-day treatment regimen initiated 14 dpi, but paxlovid-treated animals experienced rebound virus replication in the upper respiratory tract seven days after treatment end. Viral RNA was detectable 28 dpi in paxlovid-treated animals, albeit not in the molnupiravir or 4’-FlU groups, when treatment was initiated 14 dpi and continued for 14 days. Low-level virus replication continued 35 dpi in animals receiving vehicle but had ceased in all treatment groups. These data indicate that late-onset DAA therapy significantly shortens the duration of persistent virus replication in an immunocompromised host, which may have implications for clinical use of antiviral therapeutics to alleviate the risk of progression to severe disease in highly vulnerable patients.

**Importance:** Four years after the onset of the global COVID-19 pandemic, the immunocompromised are at greatest risk of developing life-threatening severe disease. However, specific treatment plans for this most vulnerable patient group have not yet been developed. Employing a CD4^+^ and CD8^+^ T cell-depleted immunocompromised mouse model of SARS-CoV-2 infection, we explored therapeutic options of persistent infections with standard-of-care paxlovid, molnupiravir, and the experimental therapeutic 4’-FlU. Late-onset treatment initiated 14 days after infection was efficacious, but only 4’-FlU was rapidly sterilizing. No treatment-experienced viral variants with reduced susceptibility to the drugs emerged, albeit virus replication rebounded in animals of the paxlovid group after treatment end. This study supports the use of direct-acting antivirals for late-onset management of persistent SARS-CoV-2 infection in immunocompromised hosts. However, treatment courses likely require to be extended for maximal therapeutic benefit, calling for appropriately powered clinical trials to meet the specific needs of this patient group.

## Introduction

COVID-19 has caused approximately 800 million confirmed cases worldwide to date, resulting in over 7 million deaths and millions of hospitalizations (1). Although downgraded from pandemic to endemic status in 2023, the evolution of the etiologic agent, severe acute respiratory syndrome coronavirus 2 (SARS-CoV-2), continues, fueling recurring infection waves carried by viral variants with altered susceptibility to pre-existing antiviral immunity in the population (2, 3). Of particular concern are prolonged infection of the immunocompromised (4–6) and long COVID syndrome, which lasts for months after infection and affects approximately 7.5% to 41% of all adults in the United States (7, 8). Symptoms include fatigue, shortness of breath, fever, headaches, sleep disturbances, and cognitive impairments (8–11), which dramatically affect quality of life of the patient. Individuals with impaired immune function due to genetic condition, cancer, and other diseases, or recipients of immunosuppressive therapy, such as transplant patients, are at heightened risk of developing severe COVID-19 with increased morbidity and mortality (12–15). Currently, specific treatment plans for COVID-19 management in immunocompromised individuals are lacking, since randomized clinical trials were not powered to assess efficacy in this patient subgroup (16, 17) and different types of naturally acquired immunodeficiencies will likely require distinct pharmacological management (4, 18, 19). Antiviral and immunomodulatory drugs are therefore recommended for use at the doses and durations that are applied to the general patient population (4).

Different small animal models of SARS-CoV-2 infection have been developed, which resemble distinct presentations of human disease. However, none of the models recapitulate prolonged infection of an immunocompromised host. Ferrets and Syrian golden hamsters are highly permissive for SARS-CoV-2, supporting efficient viral replication in the upper (20–23) or upper and lower (24, 25) respiratory tract, respectively, in each case combined with efficient viral transmission to uninfected contact animals (20, 26–28). Clinical signs in either model remain mild, and infection is fully cleared within 5 to 10 days (23, 26, 29, 30). Roborovski dwarf hamsters are likewise highly susceptible to SARS-CoV-2, including the recently emerged VOC alpha to omicron (23, 26, 31). The dwarf hamster model recapitulates severe COVID-19 with viral pneumonia and acute lung tissue damage, resulting in death of 50-90% of animals, depending on the VOC studied, within 4-7 days after infection (23, 26). Standard C57BL/6 mice support replication of some VOC without prior viral adaptation (32, 33), but animals do not develop any clinical signs and rapidly clear the infection (34, 35). Although transgenic K18-hACE2 mice expressing human ACE2 are highly susceptible to SARS-CoV-2, the disease presentation does not resemble that seen in human patients and animals succumb to viral encephalitis, due to a non-physiological tissue distribution of the ACE2 receptor (36). However, a recently reported C57BL/6 mouse model of immunocompromised animals lacking CD4^+^ and CD8^+^ T cell populations claimed SARS-CoV-2 replication in the upper respiratory tract for four weeks after infection (37).

Seeking to explore therapeutic options for high-risk immunocompromised patient populations, we examined the efficacy of late-onset antiviral therapy of tandem CD4^+^ and CD8^+^ T cell-depleted mice infected with VOC beta B.1.351, comparing two drugs in clinical use, paxlovid (38) and molnupiravir (39, 40), and the experimental therapeutic 4’-FlU (21, 41, 42). Oral treatment with either of these therapeutics initiated 14 dpi and continued twice daily (b.i.d., paxlovid and molnupiravir) or once daily (q.d., 4’-FlU) provided significant benefit, but only 4’-FlU was rapidly sterilizing. Virus rebounded in the upper respiratory tract of paxlovid-treated animals 28 dpi, seven days after treatment end. These results contribute to establishing treatment paradigms for improved management of prolonged SARS-CoV-2 infection in an immunocompromised host.

## Results

### C57BL/6 CD4^+^ and CD8^+^ T cells depletion model

To assess efficacy of DAAs in immunocompromised hosts, we employed a CD4^+^ and CD8^+^ T cell-depleted C57BL/6 mouse model, infecting animals with SARS-CoV-2 VOC beta B.1.351 (37). Animals received a cocktail of anti-CD4/anti-CD8 mAbs or isotype control, followed by assessment of depletion efficiency in regular time intervals (figure 1a). Body weight changes of animals in the initial depletion phase were minor, and indistinguishable from those of animals receiving isotype control (figure 1b), suggesting that they are a consequence of frequent injection. At predefined timepoints, blood, nasal turbinates, and lungs were harvested from animal subgroups and subjected to flow cytometric analysis. The depletion regimen resulted in nearly complete and persistent ablation of CD4^+^ and CD8^+^ T cells in circulation and upper and lower respiratory tract compartments compared to animals in the isotype control groups (figure 1c-e), which is consistent with data published for this depletion regimen (37).

**Figure 1:**
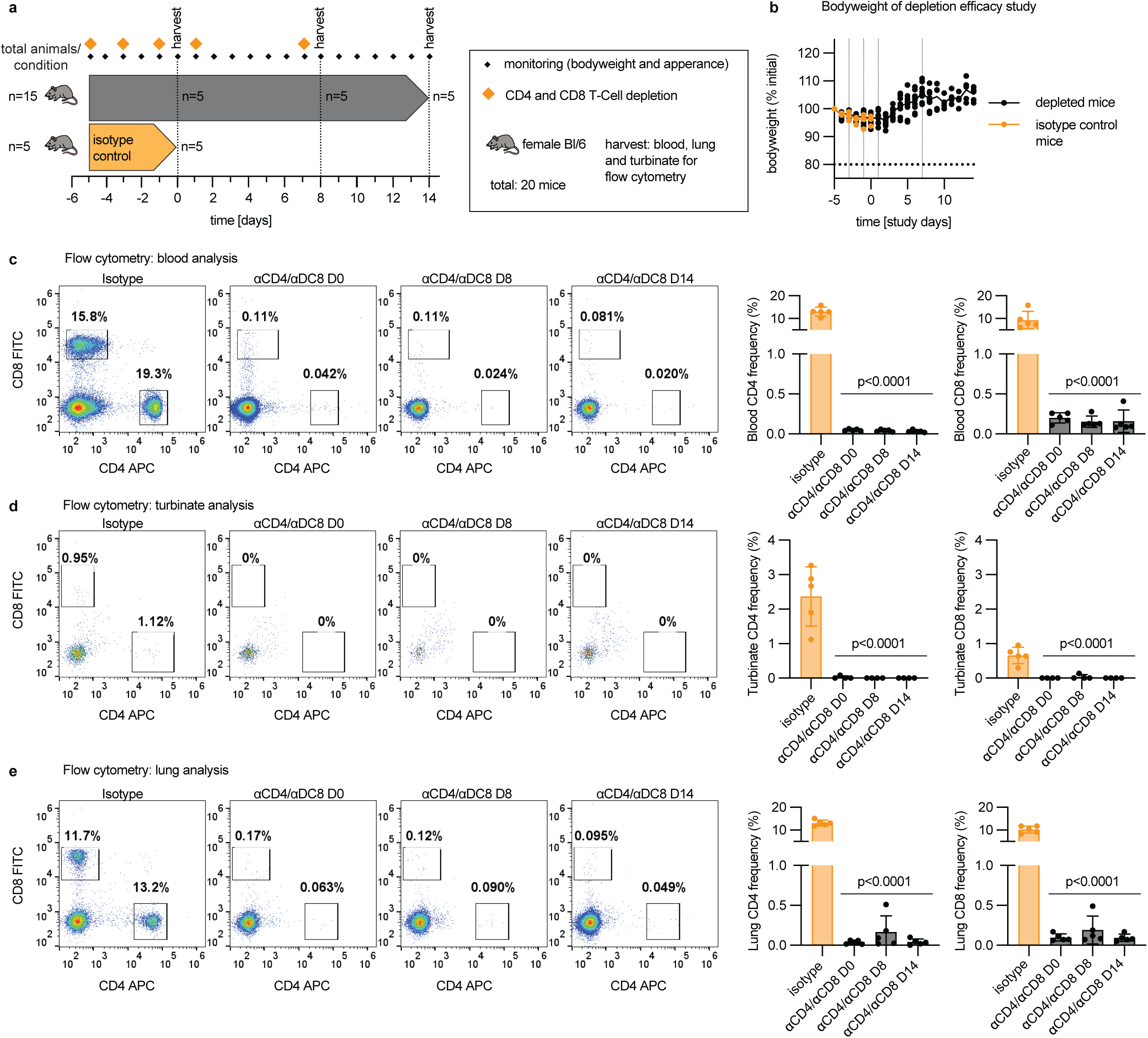
C57BL/6 CD4^+^/CD8^+^ T cell mouse depletion model. **a:** Study schematic. Blood, turbinates and lungs were harvested and processed for flow cytometry; n numbers as specified. **b:** Bodyweight curves of mice, normalized to the initial weight before the first depletion injection (study day −5). Graphs show each replicate (aligned). Horizontal dashed line denotes the predefined 20% weight-loss endpoint criteria and vertical dashed lines indicate time of depletion injections. **c-e:** Flow cytometric analysis of blood (c), turbinates (d) and lungs (e). A representative plot is shown for each analysis and condition. Right panels show CD4 and CD8 frequencies (in %). Data represent mean ± SD, p-values based on 1-way ANOVA with Dunnett post-hoc test.

### Benefit of late-onset antiviral treatment during prolonged SARS-CoV-2 infections

We selected paxlovid (250 mg/kg nirmatrelvir, 83.3 mg/kg ritonavir), molnupiravir (250 mg/kg), and 4’-FlU (EIDD-2749) (10 mg/kg) to assess efficacy of DAAs to control SARS-CoV-2 infections in immunocompromised hosts. All treatments were delivered through oral gavage, paxlovid and molnupiravir in a b.i.d. regimen, 4’-FlU was administered q.d. Mice were euthanized at pre-defined endpoints (figure 2a), and organs were collected and processed for follow-up analysis. Loss of animal bodyweight was moderate, independent of treatment, and weight stabilized 4 days after infection of mice with VOC beta B.1.351 (figure 2b). A comparison of depleted and mock-infected mice with animals not subjected to any depletion regimen, but infected with VOC beta, revealed that in this model the initial weight loss was due to a combination of stress caused by the repeated IP injection and ketamine anesthesia, rather than the result of viral pathogenesis (figure 2c-d).

**Figure 2:**
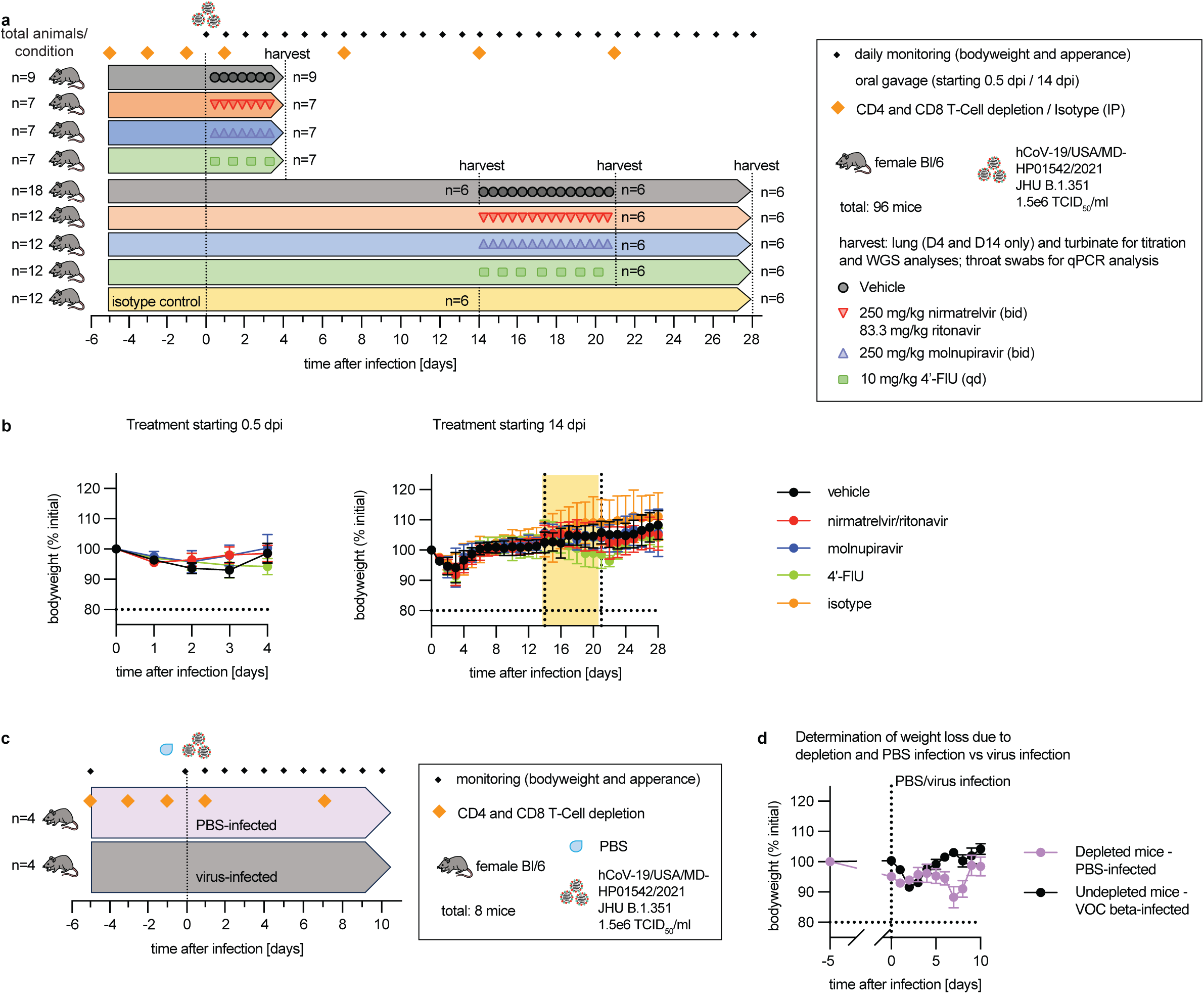
Clinical signs of immunocompromised host in acute and late-onset treatment studies. **a:** Study schematic, n numbers per condition are specified. **b:** Bodyweight measurements of different treatment regimen groups. Left panel, immediate treatment groups initiated 0.5 dpi; right panel, late onset treatment groups initiated 14 dpi. Yellow box denotes the 7-day treatment window. Symbols represent group means ± SD, lines connect means. Dashed horizontal line denotes 20% weight-loss mark. **c:** Study schematic exploring the reason for initial body weight loss. Mice were depleted and PBS mock-infected, or left undepleted and infected with hCoV-19/USA/MD-HP01542/2021 JHU B.1.351, followed by monitoring for 10 days; n numbers are specified. **d:** Bodyweight measurements. Mice were weighed daily and normalized to the initial weight before study start (day −5). Symbols represent group means ± SD, lines connect means. Horizontal dashed line denotes 20% weight-loss mark, vertical dashed line indicates the time of infection.

Treatment first initiated 12 hours after infection resulted in a statistically significant reduction in lung (figure 3a) and nasal turbinate (figure 3b) virus load 3.5 dpi in animals of the 4’-FlU group, whereas paxlovid and molnupiravir only moderately lowered lung virus titers in the immunocompromised host compared to animals of the vehicle group. Immune-competent animals had cleared infectious particles from the upper respiratory tract by 14 dpi, whereas CD4 and CD8-depleted animals maintained a robust virus load of approximately 10×10^4^ TCID_50_ units/g tissue in nasal turbinates (figure 3c). At 14 dpi, no infectious particles were detectable in the lower respiratory tract of animals in either the depletion or isotype control groups (figure 3d). Treatment initiated at that time with either drug significantly reduced virus load in turbinates by 21 dpi (figure 3e). However, infectious particles were undetectable in the molnupiravir and 4’-FlU groups 28 dpi, 7 days after treatment end, whereas paxlovid was not sterilizing (figure 3f). Quantitation of viral RNA copies in throat swabs collected 4, 14, 21, and 28 dpi confirmed these viral titration results (figure 3g).

**Figure 3:**
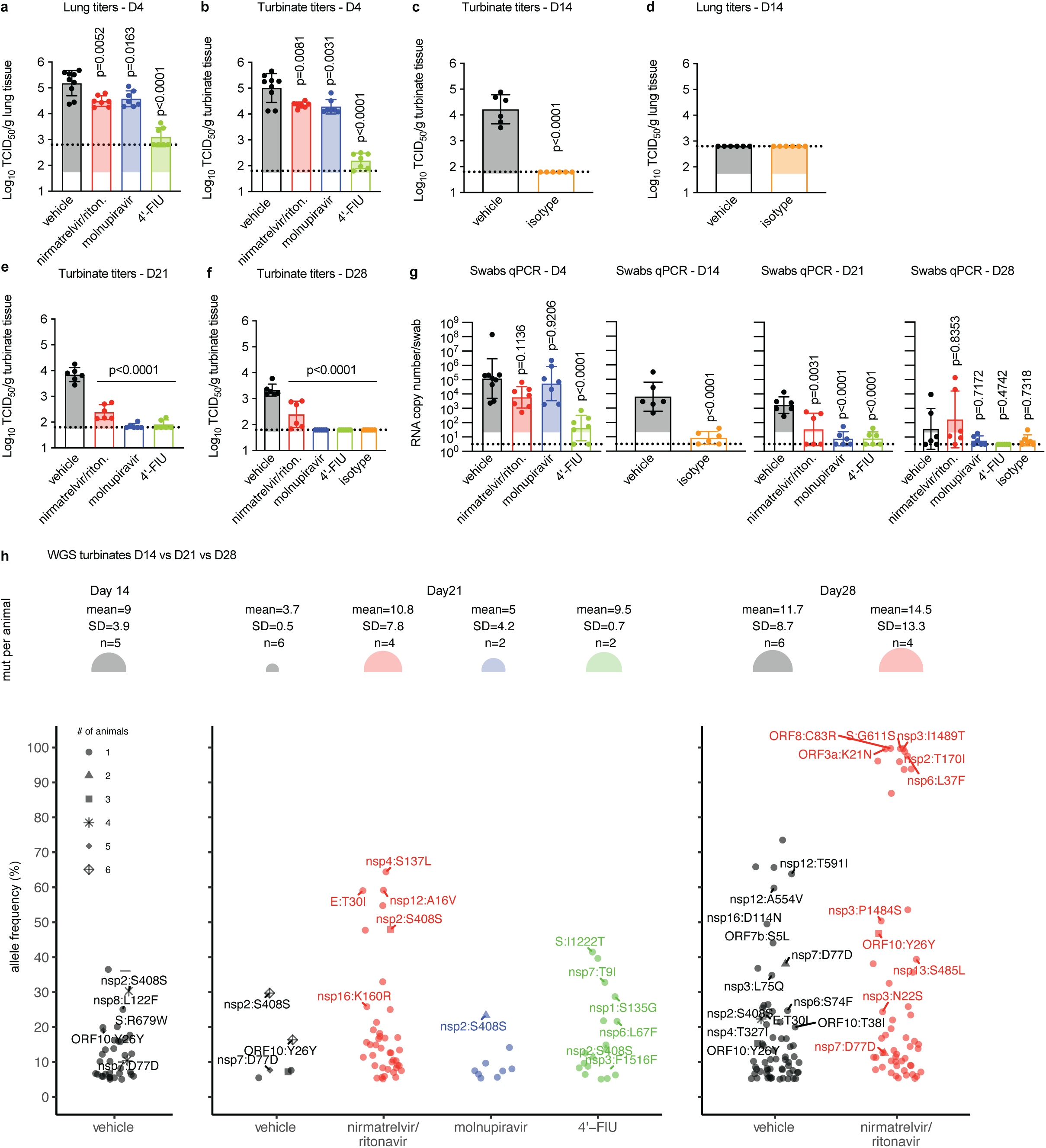
Efficacy of late-onset antiviral treatment during prolonged SARS-CoV-2 infections. **a-f:** Lung and turbinate viral titers 4 dpi (D4; a,b) and 14 dpi (D14; c,d), and turbinate titers 21 dpi (D21; e) and 28 dpi (D28; f). Columns represent geometric means ± geometric SD, symbols denote individual animals. Statistical analysis with 1-way ANOVA and Tukey’s post-hoc test (a,b,e,f) or two-tailed unpaired t-test (c); P values are shown. Dashed lines indicate limit of detection (LOD). **g:** Viral RNA copy numbers in throat swabs collected 4, 14, 21 and 28 dpi. Columns represent geometric means ± geometric SD, symbols denote individual animals. Analyses with 1-way ANOVA with Tukey’s post-hoc test or two-tailed unpaired t-test (D14 panel); P values are shown. Dashed line, LOD. **h:** Whole-genome sequencing of virus populations recovered from animals 14, 21, and 28 dpi. Top panel, differences in number of variants per animal across treatment groups and timepoints. No statistically significant differences were noted. Circle size corresponds to average number of single nucleotide variants/animal that appeared in 2 NGS technical replicates performed, with mean allele frequency ≥5% and depth ≥100. Bottom panel, unique variants identified in each treatment group plotted based on relative allele frequency. Changes in nonsynonymous mutations that emerged with ≥20% allele frequency or in more than one animal/condition are specified. Shape of symbols denotes the number of animals/condition in which each mutation was found.

Total RNA was extracted from input inoculum virus and turbinate samples of D14 (vehicle), D21 (vehicle, paxlovid, molnupiravir and 10 mg/kg 4’-FlU) and D28 (vehicle and paxlovid) and analyzed via WGS. We first examined overall mutation accumulation across the viral genomes recovered. The mean number of viral mutations per animal relative to the inoculum genome was not statistically significant across treatment groups after 7 days of treatment (21 dpi), across the three timepoints for vehicle-treated animals (14, 21, and 28 dpi), nor across the two timepoints available for paxlovid-treated animals (21 and 28 dpi, figure 3h). In contrast to other animal species infected experimentally or naturally with SARS-CoV-2 (20, 43), no longitudinal mouse-characteristic adaptations emerged during persistent infection of the immunocompromised mice.

We next examined viral sequences for mutations that arose in specific treatment groups with higher allele frequencies. No fixed mutations were recovered in any of the viruses after seven days of treatment (21 dpi). However, viruses from two 4’-FlU-treated animals developed mutations relative to the inoculum at 21 dpi with >20% allele frequency: nsp1:S135G, S:I1222T and nsp7:T9I in animal 1, and nsp6:L67F in animal 2. These mutations are uncommon in the GISAID database, only identified in 0.004-0.008% of genomes to date. Viruses from three paxlovid-treated animals developed mutations relative to the inoculum at 21 dpi with >20% allele frequency; nsp16:K160R and nsp4:S137L in animal 1, E:T30I in animal 2, and nsp12:A16V in animal 4. Remarkably, Nsp16:K160R is a canonical Omicron XBB.1.9 mutation present in 1.3% of GISAID genomes. Nsp4:S137L is a canonical beta variant mutation (44), though not present in the inoculum used in this study, and is present in 0.227% of GISAID genomes. E:T30I is relatively uncommon, represented in 0.010% of GISAID genomes, but has been previously associated with persistent infections in humans (45). E:T30I was also observed in one vehicle-treated animal at 14 dpi (13% allele frequency), and one vehicle-treated animal at 28 dpi (24% allele frequency). Nsp12:A16V was identified in a remdesivir-resistance study, but did not change viral susceptibility to the drug and is present in 0.073% of GISAID genomes. Virus populations from two animals in the paxlovid-experienced group developed fixed alternative alleles seven days after the end of treatment (28 dpi); nsp6:L37F and nsp2:T170I in animal 2, and ORF3a:K21N, nsp3:I1489T, S:G611S, and ORF8:C83R in animal 4. Nsp6:L37F has been linked to asymptomatic infections (46) and has been identified in 1.7% of GISAID sequences. The other fixed alleles are relatively uncommon among GISAID genomes.

### Drug exposure in mouse plasma and tissues

To probe for a link between differential drug efficacy and exposure, we determined plasma and tissue concentrations in a single-dose oral PK study (figure 4a). Nirmatrelvir exposure in plasma was approximately one order of magnitude higher than that of N^4^-hydroxycytidine, the circulating metabolite of molnupiravir (47, 48), and approximately four-fold higher than 4’-FlU exposure, respectively, in the 12-hour period after dosing (figure 4b). These differences in drug plasma levels were not translated to higher nirmatrelvir tissue exposure at approximate peak (3 hours after dosing) and trough (12 hours after dosing for the b.i.d. regimen) compared to the bioactive triphosphate anabolite form of molnupiravir, N^4^-hydroxycytidine-TP (figure 4c). In contrast, the bioactive anabolite of 4’-FlU, 4’-FlU-TP, showed prolonged presence in all tissues analyzed 24 hours after dosing (trough for the q.d. regimen). Of the tissues analyzed, nirmatrelvir exposure at trough was highest in nasal turbinates. Due to high sensitivity of triphosphates to hydrolysis, levels of N^4^-hydroxycytidine-TP and 4’-FlU-TP in nasal turbinates could not be determined.

**Figure 4:**
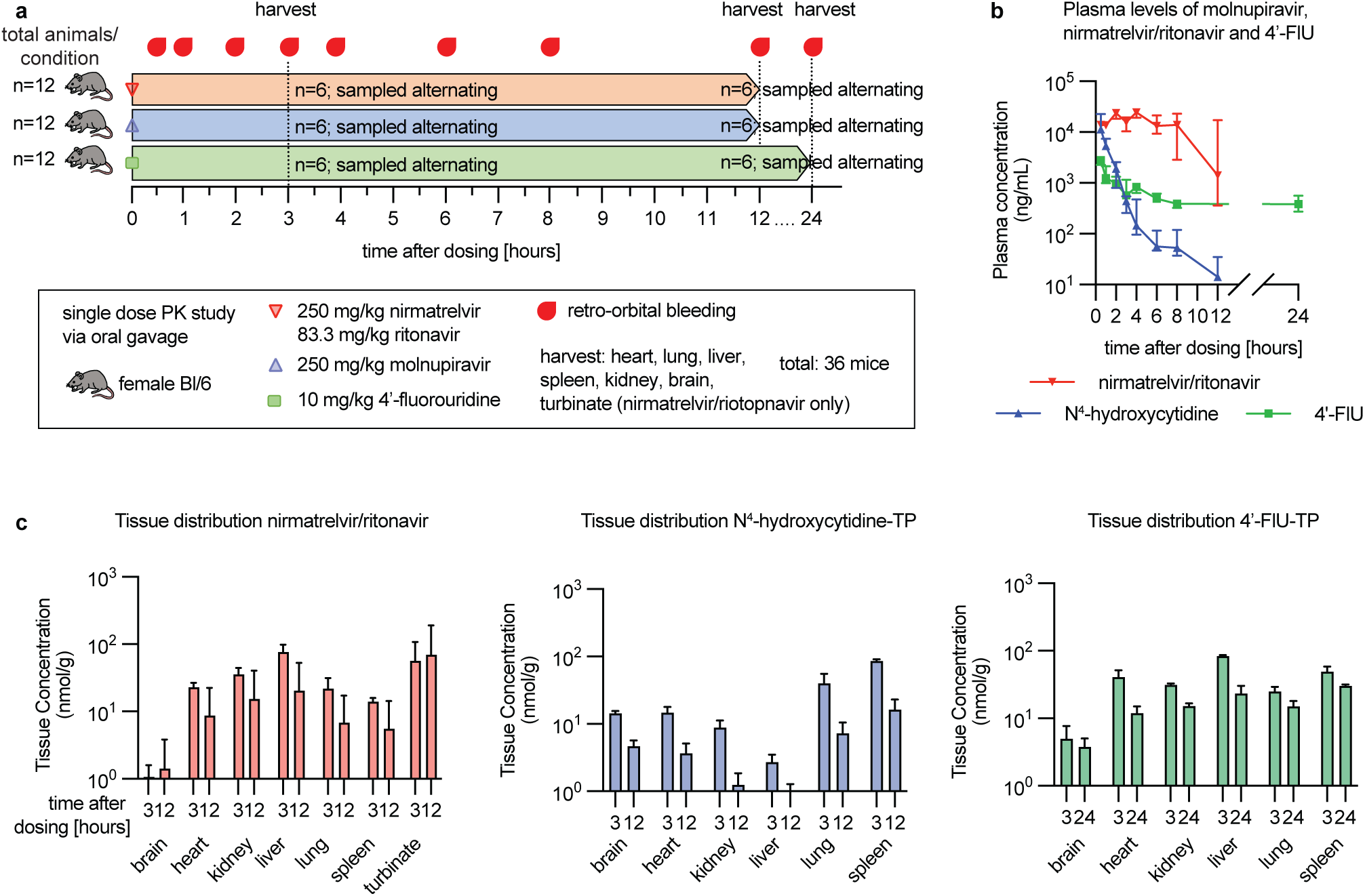
Drug exposure in mouse plasma and tissues after single oral dose. **a:** PK study schematic. Mice were dosed and bled as indicated, n-numbers are specified. **b:** Plasma concentrations of all drugs tested, symbols represent group medians ± 95% confidence intervals (CI), lines connect medians. **c:** Drug tissue distribution in selected organs 3 and 12 hours (nirmatrelvir/ritonavir, molnupiravir), or 24 (4’-FlU) hours, after dosing. For molnupiravir and 4’-FlU, the bioactive triphosphate (TP) forms of molnupiravir and 4’-FlU, N^4^-hydroxycitidine-TP and 4’-FlU-TP, respectively, are shown. Columns represent group means ± SD.

### Effect of prolonged treatment on persistent virus replication

To explore whether a longer treatment course prevents virus rebound in the nirmatrelvir/ritonavir group, we dosed animals for two consecutive weeks from 14 dpi to 28 dpi and assessed virus load in the upper respiratory tract 28 and 35 dpi (figure 5a). At treatment initiation, robust virus replication was again observed in the upper respiratory tract of CD4 and CD8-depleted animals, but not in the isotype control group (figure 5b), which was consistent with high abundance of viral RNA in throat swabs sampled from the depletion group (figure 5c). Prolonged treatment was sterilizing by 28 dpi in all treatment arms (figure 5d) and virus replication did not rebound in the upper respiratory tract of these animals 7 days after treatment end (figure 5e). However, viral RNA copy numbers in throat swabs of paxlovid-treated animals were similar to those present in the vehicle group, whereas essentially no viral RNA was detectable in the molnupiravir and 4’-FlU groups or in immune-competent animals that had received isotype control (figure 5f). By 35 dpi, viral RNA was undetectable in nearly all animals, including those of the vehicle-treated group (figure 5g).

**Figure 5:**
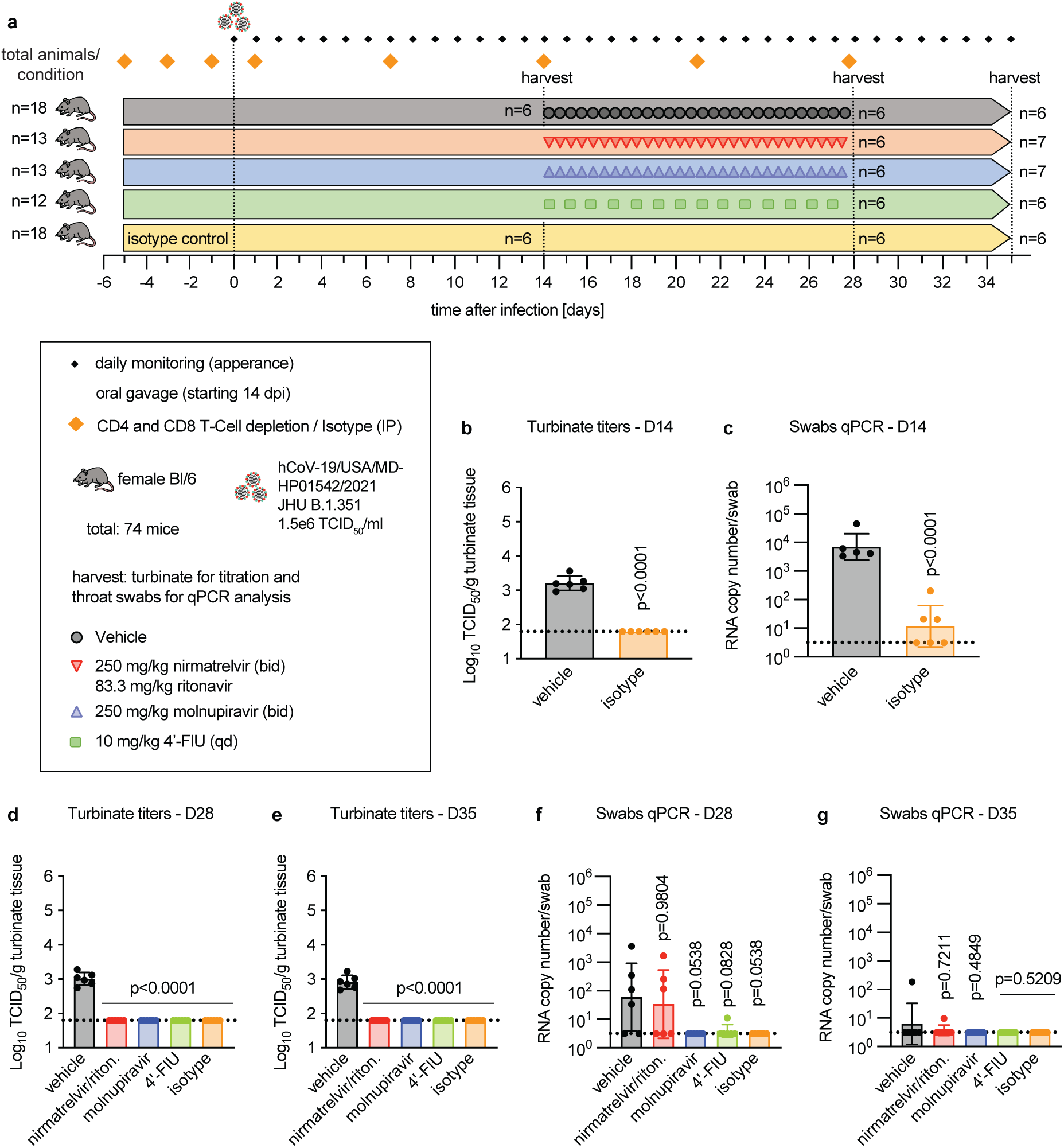
Effect of prolonged treatment on controlling persistent virus replication. **a:** Study schematic, n-numbers are specified. **b,c:** Turbinate titers (b) and viral RNA copies in throat swabs (c) of vehicle-treated depleted or isotype-control animals 14 dpi (D14); dashed line denotes LOD. **d-g:** Turbinate titers and viral RNA copies in throat swabs determined 28 dpi (d,f) and 35 dpi (e,g) in immunocompromised animals of the different treatment groups or isotype control; dashed line denotes LOD. Columns (b-g) represent geometric means ± geometric SD, symbols denote individual animals. Analysis with unpaired two-tailed t-test (b,c) or 1-way ANOVA with Tukey’s post-hoc test (d-g); P values are shown.

## Discussion

Using a mouse model of prolonged SARS-CoV-2 replication in immune-compromised hosts, we examined the efficacy of late-onset treatment with two DAAs currently in clinical use and an advanced-stage experimental therapeutic to shorten the duration of persistent virus replication. Our results support three major conclusions:

Firstly, the hallmark notion that therapeutic time windows of DAAs used against rapidly replicating RNA viruses are narrow (49, 50) does not necessarily extend to an immunocompromised host. Each of the three antivirals tested in this study statistically significantly reduced viral burden in the upper respiratory tract by several orders of magnitude when treatment was first initiated 14 dpi, when immune-competent hosts had fully cleared the infection. If equally applicable to other host species including humans, these results support the use of DAAs for mitigation of persistent SARS-CoV-2 infection in highly vulnerable immunocompromised patients. Unlike high-risk human patients, however, immunocompromised mice infected with SARS-CoV-2 did not develop viremia, major clinical signs, or progressed to severe, potentially life-threatening pneumonia. Since clinical complications arise from immunopathogenesis following the human antiviral response to SARS-CoV-2 (51–53), which is not equally pronounced in mice, the model may not fully recapitulate the therapeutic benefit that can be expected from late-onset DAA administration to immunocompromised hosts of other species.

Secondly, late-onset treatment with all three therapeutics significantly reduced virus burden compared to the vehicle group, However, only molnupiravir and 4’-FlU were sterilizing after a 7-day regimen. In contrast, low-level virus replication continued in paxlovid recipients, possibly creating opportunity for virus rebound after treatment end and/or continued virus shedding from immunocompromised hosts. In previous work, we noted limited efficacy of paxlovid in controlling SARS-CoV-2 replication in the ferret upper respiratory tract, which we attributed to low nirmatrelvir exposure in ferret nasal turbinates compared to other soft tissues (23). However, the PK analysis carried out in this study revealed that distinct tissue exposure is species-specific, since we detected high and sustained nirmatrelvir levels in mouse turbinates, which could not be reconciled with persistent virus replication in the upper respiratory tract. Despite lower exposure of N^4^-hydroxycytidine-TP in mouse tissues than in other host species (23, 54, 55), molnupiravir provided better control of virus replication in the upper respiratory tract, which could reflect the distinct mechanisms of action of the two drugs, viral mutagenesis induced by molnupiravir (41, 56–58) versus M^pro^ inhibition by nirmatrelvir (23, 41, 59), and is consistent with a slightly lower rate of viral rebound in human patients after a molnupiravir versus paxlovid treatment course (60). Consistent with overall superior tissue exposure of its bioactive triphosphate form, 4’-FlU most effectively controlled virus replication when first administered in the acute or persistent infection phase. These results underscore strong antiviral potency of 4’-FlU against SARS-CoV-2 in ferrets (21) and unrelated viral pathogens such as respiratory syncytial virus (21) and influenza viruses (42) in different animal model systems, emphasizing the high developmental potential of the compound.

Thirdly, despite prolonged viral replication in the presence of inhibitor and replication rebound in the paxlovid group after treatment end, whole genome sequencing of virus populations recovered from the animals did not reveal the emergence of any subpopulations with reduced susceptibility to inhibition. However, one variant previously associated with persistent infection, E:T30I, emerged in a nirmatrelvir/ritonavir-treated animal and two vehicle-treated animals, which supports the physiological relevance of this animal model. Although resistance to paxlovid has been induced experimentally (61, 62) and some viral alleles associated with reduced viral susceptibility to nirmatrelvir have been identified in circulating SARS-CoV-2 populations (63, 64), widespread viral resistance to paxlovid has not yet emerged as a clinical problem. Our results in the immunocompromised mouse model support this conclusion and suggest a relatively high genetic barrier to viral escape *in vivo*. Though some minor alleles were identified in 4’-FlU-treated animals, effective suppression of SARS-CoV-2 replication by molnupiravir and 4’-FlU made *in vivo* viral adaptation to the drugs even less feasible. Moreover, no confirmed resistance mutations to molnupiravir have been identified experimentally (65) or emerged in clinical studies (66). We have previously reduced susceptibility-profiled 4’-FlU against influenza viruses and found that resistance was moderate, resulted in viral attenuation and/or reduced transmissibility, and could in all cases be overcome *in vivo* by administration of standard or slightly higher dose levels (67). Taken together, the mouse model provides no indication of a heightened rate of viral escape in drug-treated immunocompromised hosts with prolonged viral replication phase.

This study supports the use of DAAs for the treatment of persistent SARS-CoV-2 infection in immunocompromised hosts. However, the mouse model suggests that, depending on the drug used, treatment courses in immunocompromised individuals require to be adjusted compared to those established for the general patient population for maximal efficacy, calling for appropriately powered randomized clinical trials to meet the specific needs of this patient subgroup.

## Material and Methods

### Study design

Cells and mice were used as *in vitro* and *in vivo* models to examine the potency of nirmatrelvir/ritonavir, molnupiravir, and 4’-FlU in a prolonged SARS-CoV-2 infection model. Viruses were administered through intranasal inoculation and virus load monitored periodically in respiratory tissues of mice extracted 4 days, 14 days, 21 days, 28 days, and 35 days after infection. Virus titers were determined through TCID_50_-titration and RNA copy numbers via qPCR in thoracal swabs.

### Virus stock growth

Vero E6-TMPRSS2 cells were infected with hCoV-19/USA/MD-HP0142/2021 JHU B.1.351 when reaching 80% confluency at a multiplicity of infection (MOI) of 0.001 TCID_50_ units/cell.

Cells were observed for cytopathic effect (CPE), and virus was harvested after 2.75 days. Virus was spun down for 10 minutes at 1000 rpm and 4°C, and cleared supernatant was aliquoted and frozen at −80°C before titering via TCID_50_.

### TCID_50_ titration

VeroE6-TMPRSS2 cells were seeded in 96-well plates and incubated overnight. Viral samples were diluted 1:10 and transferred to the cells. Plates were scored 3 days after infection via light microscopy (Reed and Muench method).

### Flow cytometry studies

Female C57BL/6 mice (Jax), 6 weeks of age were rested for 1 week before study start. Mice (n=5) were depleted with anti-mouse CD4 (Fisher Scientific 50561964) and anti-mouse CD8 (Fisher Scientific 50562192) mAb or isotype control (IP, 100 µl, 200 µg each) at predefined timepoints. Upon harvesting, the left cranial lung lobe was dissected from mice and minced with scissors before digestion for 30 minutes at 37 °C with RPMI containing 5% FBS, DNase, and Collagenase type IV. Lungs were further homogenized via swishing with a 5 ml syringe every 10 minutes during digestion. Digestion was followed by passage through a 70 μm cell strainer to remove tissue fragments. Red blood cells were lysed with red blood cell lysis buffer (BioLegend). Lung cells were re-suspended in 1 ml of RPMI containing 5% FBS. 100 µl of blood was collected into a 1.5 ml Eppendorf tube containing 100 μl sodium citrate (to prevent blood clotting) using the retro-orbital bleeding method. Blood was centrifuged at 5000 rpm for 15 minutes at 4 °C and serum discarded. Cells were resuspended in 5 ml of red blood cell lysis buffer and incubated on ice for 5 minutes before removing the lysed red blood cells via centrifugation. Blood leukocytes were then washed in 15 ml of PBS and re-suspended in 200 μl of RPMI containing 5% FBS. Nasal turbinates were harvested, and a single-cell suspension was obtained by crushing the turbinate through a 70 µm cell strainer, centrifuged, and re-suspended in 200 μl of RPMI containing 5% FBS. For flow cytometry analysis, the prepared cells were blocked with 2.4G2 (anti-CD16/anti-CD32) at a concentration of 1 μg/million cells in 200 μl PBS for 15 minutes at 4°C, followed by 1 wash with PBS to remove all residue of 2.4G2. Cells were incubated with a cocktail of conjugated monoclonal antibodies of CD45 BV605, CD8a FITC, and CD4 APC at 4°C for 30 minutes. Multi-parameter analysis was performed on a CytoFlex (Beckman Coulter) and analyzed via FlowJo software (Tree Star).

### Mouse PK studies

Twelve mice for each compound were gavaged orally with either 250 mg/kg nirmatrelvir and 83.3 mg/kg ritonavir (paxlovid), 250 mg/kg molnupiravir, or 10 mg/kg 4’-FlU, respectively. Blood was collected retro-orbitally 30 minutes, 1 hour, 2 hours, 3 hours, 4 hours, 6 hours, 8 hours, and 12 hours (molnupiravir and paxlovid) or 24 hours (4’-FlU) after dosing. Blood was spun for 10 minutes at 2500 rpm and 4°C and plasma aliquoted and stored at −80°C. Organs (lung, heart, liver, spleen, kidney, brain, and turbinates (paxlovid only)) were harvested 3 hours and 12 hours or 24 hours after dosing, respectively. Organs were flash-frozen in liquid nitrogen before transfer to −80°C.

### LC/MS/MS

Aliquots of mouse plasma were extracted with acetonitrile including appropriate internal standards. Samples of frozen animal tissue were extracted with cold (4°C) 70% acetonitrile in water including internal standards by homogenization in a Lysera bead mill outfitted with a cryo-cooling unit (Biotage, Uppsala, Sweden). HPLC separation was performed on Agilent 1200 or 1260 systems (Agilent Technologies, Santa Clara, CA, USA) utilizing a Zorbax XDB-C18 (2.1×50 mm, 3.5μm) column (Agilent Technologies), a BEH Z-HILIC PEEK-coated (50 x 4.6 mm, 5 µm particle size) column (Waters, Milford, MA, USA), a SeQuant ZIC-pHILIC PEEK-coated (100×4.6 mm, 5 µm, PEEK coated) column (Merck Millipore, Burlington, MA, USA), or a SeQuant ZIC-cHILIC (100×4.6mm, 3 µm, PEEK coated) column (Millipore Sigma). Mass spectrometry analysis was performed on a Triple Quadrupole 5500, QTrap 7500 or Triple Quadrupole 7500 Mass Spectrometer (Sciex, Framingham, MA, USA) using electrospray ionization in multiple reaction monitoring mode or in MS/MS/MS mode. Data analysis was performed using Analyst or SciexOS software (Sciex). PK parameters were calculated using Phoenix WinNonLin 8.3 (Build 8.3.5.340; Certara, Princeton, NJ) using the non-compartmental analysis tool.

### Mouse infection studies

Female C57BL/6 mice (Jax), 6 weeks of age, were for 1 week prior to study start. Five, three and one days before SARS-CoV-2 infection and one, seven, 14, 21, and 28 days after infection, mice were depleted with anti-mouse CD4 (Fisher Scientific 50561964) and anti-mouse CD8 (Fisher Scientific 50562192) mAb or isotype control (IP, 100 µl each, 200 µg). Mice were anesthetized using ketamine and infected with 1.5×10^6^ TCID_50_ units of hCoV-19/USA/MD-HP0142/2021 JHU B.1.351, or PBS for mock infections. Animals were treated via oral gavage with either 250 mg/kg nirmatrelvir and 83.3 mg/kg ritonavir (b.i.d.), 250 mg/kg molnupiravir (b.i.d.), or 10 mg/kg 4’-FlU (q.d.) and euthanized on pre-defined endpoints on days 4, 14, 21, 28, and 35. Throat swabs (for qPCR), lungs (days 4 and 14 only, for titration via TCID_50_), and turbinates (for titration via TCID_50_ and WGS analyzes) were collected. Tissue samples were weighed, homogenized using a bead-blaster (3×30 seconds at 4°C with 30 seconds rest between cycles), spun, and aliquoted. Aliquots were kept at −80°C upon titration on VeroE6-TMPRSS2 cells. In addition, throat swabs and turbinate homogenates were processed immediately in viral RNA buffer (Zymo Research) and RLT lysis buffer (QIAGEN) and RNA was extracted following the manufacturers instructions (Zymo Quick-RNA^TM^ Viral Kit or QIAGEN RNeasy Mini Kit, respectively).

### Quantitative real time PCR

Viral RNA copy numbers in throat swabs samples were determined using and Applied Biosystems QuantStudio 3 instrument, Taqman Fast 1 Step mix (Thermo), and IP2 primers (68). Normalized RNA copy numbers per swab were calculated.

### SARS-CoV-2 whole genome sequencing

RNA from turbinate samples were sequenced in duplicate by reverse-transcription and amplification using the IDT-Swift Biosciences SARS-CoV-2 assay with v2 primers and sequenced on an Illumina NextSeq 2000 (Illumina, Inc). Reads were processed and inoculum consensus genome was generated with TAYLOR (https://github.com/greninger-lab/covid_swift_pipeline), which uses Trimmomatic for quality/adapter trimming, bbmap for alignment to the Wuhan-Hu-1 reference (Genbank accession NC_045512.2), and primerclip to remove primer sequences. The SARS-CoV-2 reads were aligned to the inoculum reference genome and variants were called with RAVA (https://github.com/greninger-lab/lava/tree/Rava_Slippage-Patch) and the inoculum consensus genome, which uses bwa-mem for alignment and VarScan2 for vcf-generation. A custom R script was used to further filter the variants and generate the figures (https://github.com/greninger-lab/sars-cov-2_mouse_model_supp). Two technical replicates of viral WGS were performed per sample, and variants that appears with ≥1% frequency and without strand-bias (<90% alternate alleles from one strand) in both technical replicates, and had mean allele frequency ≥5%, and mean depth ≥100 between technical replicates were considered for downstream analysis.

### Statistical analysis

For statistical analysis of more than two study groups, 1-way analysis of variance (ANOVA) with appropriate multiple comparison post-hoc tests as specified in figure legends was used to assess statistical difference between samples. Statistical analyses were carried out in Prism version 10.0.3 (GraphPad), and graphs were assembled in Adobe illustrator. Cartoons and study design overviews were partially generated in Biorender. The number of individual biological replicates (n values) and exact P values are shown in the schematics and figures. The threshold of statistical significance (α) was set to 0.05.

### Ethical compliance

All animal work was performed in compliance with the *Guide for the Care and Use of Laboratory Animals* of the National Institutes of Health and the Animal Welfare Act Code of Federal Regulations. Experiments involving mice were approved by the Georgia State University Institutional Animal Care and Use Committee (IACUC) under protocol A22037. All experiments using infectious material were approved by the Georgia State University Institutional Biosafety Committee (IBC) under protocol B23016 and performed in BSL-3/ABSL-3 high containment facilities of Georgia State University.

## Data availability

All raw sequencing data and inoculum consensus genome sequences were deposited under BioProject PRJNA1108751. All numerical raw data and statistical analyses are available in the supplementary Source data and Statistics files, respectively.

## Acknowledgements

We thank MK Andrews, AA Cruz, and RE Krueger for technical assistance with mass-spectrometry analyses, M Kar for providing virus stocks, and the Georgia State University Department of Animal Resources and the High Containment Core for assistance. This work was supported, in part, by public health service grants AI171403 (to RKP) and AI141222 (to RKP).

